# Subtype-specific roles of ellipsoid body ring neurons in sleep regulation in *Drosophila*

**DOI:** 10.1101/2022.09.11.507483

**Authors:** Wei Yan, Hai Lin, Junwei Yu, Timothy D. Wiggin, Litao Wu, Zhiqiang Meng, Chang Liu, Leslie C. Griffith

## Abstract

The ellipsoid body (EB) is a major structure of the central complex of the *Drosophila melanogaster* brain. 22 subtypes of EB ring neurons have been identified based on anatomical and morphological characteristics with light-level microscopy and EM connectomics. A few studies have associated ring neurons with the regulation of sleep homeostasis and structure. However, cell type-specific and population interactions in the regulation of sleep remain unclear. Employing a unbiased thermogenetic screen of collected EB drivers, we found: 1) multiple ring neurons are involved in the modulation of amount of sleep and structure in a synergistic manner; 2) analysis of data for ΔP(doze)/ ΔP(wake) using a mixed Gaussian model detected 5 clusters of GAL4 drivers which had similar effects on sleep pressure and/or depth: lines driving arousal contained R4m neurons, whereas lines that increased sleep pressure had R3m cells; 3) a general linear model analysis correlating ring cell subtype and activity-dependent changes in sleep parameters across all the lines identified several cell types significantly associated with specific sleep effects: R3p for daytime sleep promotion, and R4m for nighttime wake-promoting; and 4) another subclass, R3d cells present in 5HT7-GAL4+ neurons and in GAL4 lines from our screen which exclusively affect sleep structure, were found to contribute to fragmentation of sleep during both day and night. Thus, multiple subtypes of ring neurons distinctively control sleep amount and/or structure, and the unique highly interconnected structure of the EB and its connections with other regions of brain suggest a local-network model worth future investigation.

**SIGNIFICANCE STATEMENT:** How multiple brain regions, with many cell types, can coherently regulate sleep remains unclear, but identification of cell type-specific roles can generate opportunities for understanding the principles of integration and cooperation. The ellipsoid body (EB) of the fly brain exhibits a high level of connectivity and functional heterogeneity yet is able to tune multiple behaviours in real-time, including sleep. Leveraging the powerful genetic tools available in *Drosophila* and recent progress in the characterization of the morphology and connectivity of EB ring neurons, we identify several EB subtypes specifically associated with distinct aspects of sleep. Our findings will aid in revealing the rules of coding and integration in the brain.

## INTRODUCTION

Sleep plays critical roles in many physiological functions. Sleep regulation in the brain is a complex process modulated at the molecular, cellular, circuit and network levels (John et al., 2016; Scammell et al., 2017; Bringmann, 2018; Herice and Sakata, 2019; Liu and Dan, 2019). Previous studies in *Drosophila melanogaster* have revealed multiple cell types and neural circuits that participate in the regulation of sleep amount, structure and homeostasis.

The ellipsoid body (EB) contributes to regulation of multiple behaviors including spatial orientation, navigation, arousal and sleep (Bausenwein et al., 1994; Lebestky et al., 2009; Ofstad et al., 2011; Seelig and Jayaraman, 2015; Fisher et al., 2019; Kim et al., 2019; Kottler et al., 2019). As one of the central structures on the midline of the fly brain, the EB receives direct input from, and sends output to, many brain regions. This high level of connectivity positions the EB to be a center for integration of multiple information streams, including visual, motor, mechanosensory and circadian input, allowing it to functionally tune complex behaviors (Franconville et al., 2018).

The organization within the EB also exhibits complexity. With recent progress on morphology and connectivity of the EB, 22 distinct subtypes of ring neurons have been identified (Hulse et al., 2021). Each subtype of ring neuron typically contains a dendritic arborization lateral to the EB, then projects a single axon into the concentric laminated structure within the EB neuropil. The projections from each subtype of ring neuron form distinct layers within the neuropil, terminating in different rings at specific depths along the anterior-posterior axis where they interconnect (Hanesch et al., 1989; Young and Armstrong, 2010; Lin et al., 2013). These connections, between neurons of the same type, provide each ring neuron’s strongest inputs (Isaacman-Beck et al., 2020; Hulse et al., 2021) and suggest a structural basis for local communication and synergism for sleep regulation.

In spite of the growing understanding of EB connectivity, specific roles for each subtype of ring neuron in sleep is limited. One subtype of R5 neuron (initially referred to as R2) has been shown to drive a persistent sleep upon secession of thermoactivation, suggesting a role in sleep drive and homeostasis (Donlea et al., 2014; Liu et al., 2016; Pimentel et al., 2016). Another study showed that single R5 neurons get synchronized by circadian input and the power of slow-wave oscillations in R5 neurons has been associated with increased sleep drive (Raccuglia et al., 2019). 5HT7-GAL4+ EB neurons, which consist of several subtypes and are modulated by serotonergic signaling, can regulate sleep architecture (Liu et al., 2019). In spite of these important findings, the scope of ring neuron involvement in the regulation of sleep is not clear.

In the present study, we take an unbiased approach, screening 34 drivers that label different combinations of subtypes of ring neurons by thermoactivation using the warmth-sensitive cation channel dTrpA1 (Hamada et al., 2008). Most drivers label multiple ring neurons, and activation of many drivers resulted in significant changes in sleep amount and/or sleep structure. The complexity of the tools and phenotypes necessitated developing computational approaches for assessing the importance of each subtype. Using P(wake) and P(doze) analysis with a mixed Gaussian model, 5 clusters of drivers were found to regulate sleep depth and pressure during the day and/or at night, respectively. Furthermore, a general linear model analysis based on the GAL4 expression pattern and the sleep behavior upon 24-hour activation suggest several types of ring neuron contribute to sleep regulation consistent with and extending the findings from the Gaussian model. Finally, using genetic suppression of intersected population strategy, we identified a subpopulation of neurons which is sufficient to fragment sleep during both day and night. Although how the ring neurons cooperate to coherently modulate sleep is not yet clear, the identification of roles for specific cell types provides an important piece of the puzzle.

## MATERIALS AND METHODS

### EXPERIMENTAL MODEL AND SUBJECT DETAILS

#### Animals

Unless specified, flies were reared on standard cornmeal food (Each 1 liter H_2_O: 70g cornmeal, 50g sucrose, 10g soybean powder, 20g yeast powder, 6g agar, and 3g Methyl 4-hydroxybenzoate) at 23°C with 60% relative humidity and under a regime of 12-hour light/12-hour dark. Flies were allowed to freely mate after eclosion, and mated females aged 2~5 days were used for all experiments. GAL4 lines: R12B01 (48487), R15B07 (48678), R28D01 (47342), R28E01 (49457), R38B06 (49986), R38G08 (50020), R38H02 (47352), R41A08 (50108), R41F05-GAL4 (50133), R47F07 (50302), R48B10 (50352), R49E12 (38693), R53F11 (50443), R53G11 (69747), R54B05 (69148), R56C09 (39145), R64H04 (39323), R70B04 (39513), R70B05 (47721), R73A06 (39805), R73B05 (48312), R81F01 (40120), R84H09 (47803), Aph^c507^ (30840), C232 (30828), and R44D11-LexA (41264), UAS-dTrpAl (26263), UAS-mCD8::GFP (5136), UAS-mCD8::RFP, LexAop2-mCD8::GFP (32229), LexAop-Gal80 (32213) were ordered from the Bloomington Drosophila Stock Center. GAL4 lines of VT038828 (v201975), VT040539 (v204084), and VTO59775 (v201924) were ordered from Vienna Drosophila Resource Center (VDRC). GAL4 lines: VT012446, VT026841, VT042577, VT042759, VT045108 and VT057257 were ordered from VDRC originally, but unfortunately not available anymore. 5HT7-GAL4 was provided by Charles Nicols’ Lab. Feb170-GAL4 was generated by Günter Korge’s Lab (Siegmund and Korge, 2001). The wild type line *w^cs^* was crossed with GAL4 and UAS parental lines as genetic controls.

#### Method details

##### Sleep assay and calculation of sleep changes

F1 generation of flies were all maintained on standard food at 23 °C. 2-5 days old mated F1 female flies were individually placed into a 65 mm × 5 mm glass tube containing food (2% agar and 5% sucrose). After loading to the DAM2 system (Drosophila Activity Monitor) (Trikinetics, Waltham) at 21 °C in 12 hr: 12 hr light/dark (LD) cycles, flies were entrained for 2-3 days. Then one day baseline sleep, one day neural activation sleep as well as one day recovery sleep were recorded at 21 °C, 30 °C and 21 °C, respectively. Total sleep, the number of sleep episodes, and max episode length were analyzed for light and dark periods (LP and DP) separately, using MATLAB program (SCAMP2019v2) scripts.

To overview the effects upon activation of GAL4+ neurons, all genotypes were arranged in a descending order according to the changes of total sleep during the light period. Sleep changes were calculated by subtracting baseline day sleep of each genotype from its activation day. For genotypes with significant changes in sleep and/or sleep structure, three days’ sleep profiles of sleep time in 30 min were plotted. Sleep changes of the recovery day were also calculated. The significant difference was marked when the experimental group is different compared to both genetic controls.

##### Immunohistochemistry

Brains of adult flies were dissected in 10 mM ice-cold PBS and fixed for 20 min in PBS with 4% paraformaldehyde at room temperature. Brains were then washed three times for 5 min each in PBT (PBS with 0.5% Triton X-100). For GFP and RFP immunostaining, brains were incubated with primary antibodies (1:200, chicken anti-GFP, Abeam, Cat# ab13970; 1:200, mouse anti-GFP, Roche, Cat# AB_390913; 1:1000, rabbit anti-GFP, Invitrogen, Cat# A-l 1122; 1:200, rabbit anti DsRed, Takara, Cat# AB_10013483) in 10% NGS in PBT at 4 °C for two nights. After three times washes for 5 min each with PBT at room temperature, brains were incubated with secondary antibody at 4°C overnight. Second antibodies (488 goat anti-mouse, Invitrogen, Cat# A-11001; 488 goat anti-chicken, Invitrogen, Cat# A-l 1039; 488 goat anti-rabbit, Invitrogen, Cat# A-11008; 568 goat anti-rabbit, Thermo Fisher, Cat# A-11011) were all used in a ratio of 1:200. Samples were then washed three times for 5 min each in PBT at room temperature, and mounted on microscope slide in Vectashield mounting medium (Vector Laboratories, Inc. Car# H-1000). Finally, samples were imaged with Leica TCS SP5/LSM900 confocal microscope and analyzed using the open source of FIJI (Image-J) software.

##### Probability Analysis

The probability of transitioning from a sleep to an awake state (P(wake)), and from a wake state to a sleep state (P(doze)) were used power law distributions analysis as previously described (Wiggin et al., 2020). P(wake) and P(doze) were calculated identically, with calculation of 1-min bin of inactivity and activity reversed. The MATLAB scripts for analysis of P(wake)/P(doze) can be accessed in GitHub at https://github.com/Griffith-Lab/Fly_Sleep_Probability.

##### Mixed Gaussian Model Clustering

To figure out different effects of EB drivers on both sleep pressure and depth, we divided all significant subtypes of EB ring neurons into groups with similar distributions of delta P(Wake) and delta P(Doze), using mixed gaussian model clustering. The clustering analysis was conducted using the scripts of fitgmdist and cluster in Matlab. Given the small sample size of neuron subtypes (14 and 13 for daytime and nighttime, respectively), the number of cluster k was set to 3, 4, or 5 for both daytime and nighttime. We calculated the silhouette coefficients for each k value using the script of silhouette in Matlab and chose the final k value whose silhouette coefficient was the closest to one (Lecompte et al., 1986). The size of ellipse for each cluster was decided by the corresponding sigma values of its gaussian mixture distribution.

##### General linear model

To evaluate the effect of a specific anatomical subtype of ring neurons on sleep, the generalized linear model (GLM) (Generalized Linear Models. 2nd Edition, Chapman and Hall (CRC Press), http://dx.doi.org/10.1007/978-1-4899-3242-6) was used to estimate the weights and the corresponding statistical significance of all subtypes for each sleep parameter. The GLM analysis was conducted using the script of glmfit in Matlab (MathWorks, Natick, MA) to predict each sleep parameter under the combination of all subtypes of neurons. The input variable was defined as 1 or 0 for each subtype of ring neurons (Rl, R2, R3d, R3m, R3a, R3p, R3w, R4m, R4d, R5, and R6) when labeled or not labeled by each driver, respectively. And the corresponding output variable was the mean change rate of each sleep parameter of the same driver upon the activation to its baseline level (output variable value = (activation-baseline)/baseline). We chose the default parameters for the script of glmfit. According to the weight calculation for each subtype (Figure 5-1), a positive value represents positive relationship, and a negative value represents negative relationship between the subtype and the sleep parameter, respectively, when the corresponding p value <0.05.

##### Statistical analysis

Power analysis was conducted using the script of sampsizepwr in Matlab (MathWorks, Natick, MA) to calculate the power for the sample size in this study. The power analysis was based on the sleep parameters in drivers with significant differences from both control groups presented in the main figures. We selected the mean and standard deviation of control groups under the null hypothesis, and the mean value of experimental groups under the alternative hypothesis during the calculation of power values. Based on current sample size, >80% of the powers of significances of sleep parameters were greater than 0.9 (Figures 2-2, and 5-2).

Data were performed using GraphPad Prism 8. Group means were compared using one-way ANOVA followed by Bonferroni’s multiple comparison test when data were normally distributed, or Kruskal-Wallis test followed by Dunn’s multiple comparison test was employed when data failed passing normality test. All experiments were performed at least 2 replicates, and data presented in the figures were chosen from one representative replicate. To uniform the data presentation, all figures were prepared as mean ± SEM. To visualize all groups in the same figure clearer, error bars were not shown. Asterisk (*) indicates a significant difference: *p < 0.05, **p < 0.01, ***p < 0.001, ****p < 0.0001; N.S., not significant.

## RESULTS

### Thermoactivation of ring neurons changes sleep amount

To investigate the roles of ring neuron types, we collected 34 GAL4 drivers that label different populations of ring neurons and used them to drive the thermogenetic tool dTrpAl, allowing the use of elevated temperature to drive neuronal firing (Hamada et al., 2008). Animals were placed in DAM2 system tubes and entrained at 21 °C in a 12 hr: 12 hr light/dark (LD) cycle. Sleep was then recorded for 3 days: one day of baseline sleep at 21 °C, one day of neural activation sleep at 30 °C, then one day of recovery sleep at 21 °C (Figure 1A). Changes in sleep parameters for each genotype on the activation and recovery days were calculated by subtracting the baseline day value (Figure 1 A). Changes were only considered significant when the experimental group was different from both genetic controls. Changes in total daytime sleep of the 34 drivers on the activation day are arranged in descending order (Figure 1B), and changes of total nighttime sleep (Figure 1G) as well as changes in the number of episodes (Figure 1C and H), maximum episode duration (Figure 1D and I), P(doze) (Figure 1E and J), and P(wake) (Figure 1F and K) are displayed in the same order as the daytime sleep data to allow assessment of all parameter changes for each genotype. The color-coding of the histogram bars corresponds to the Gaussian clusters shown in Figure 4 and is also used to identify lines in Figures 2 and 5 and Figure 2-1 as part of particular clusters.

**Figure 1.**
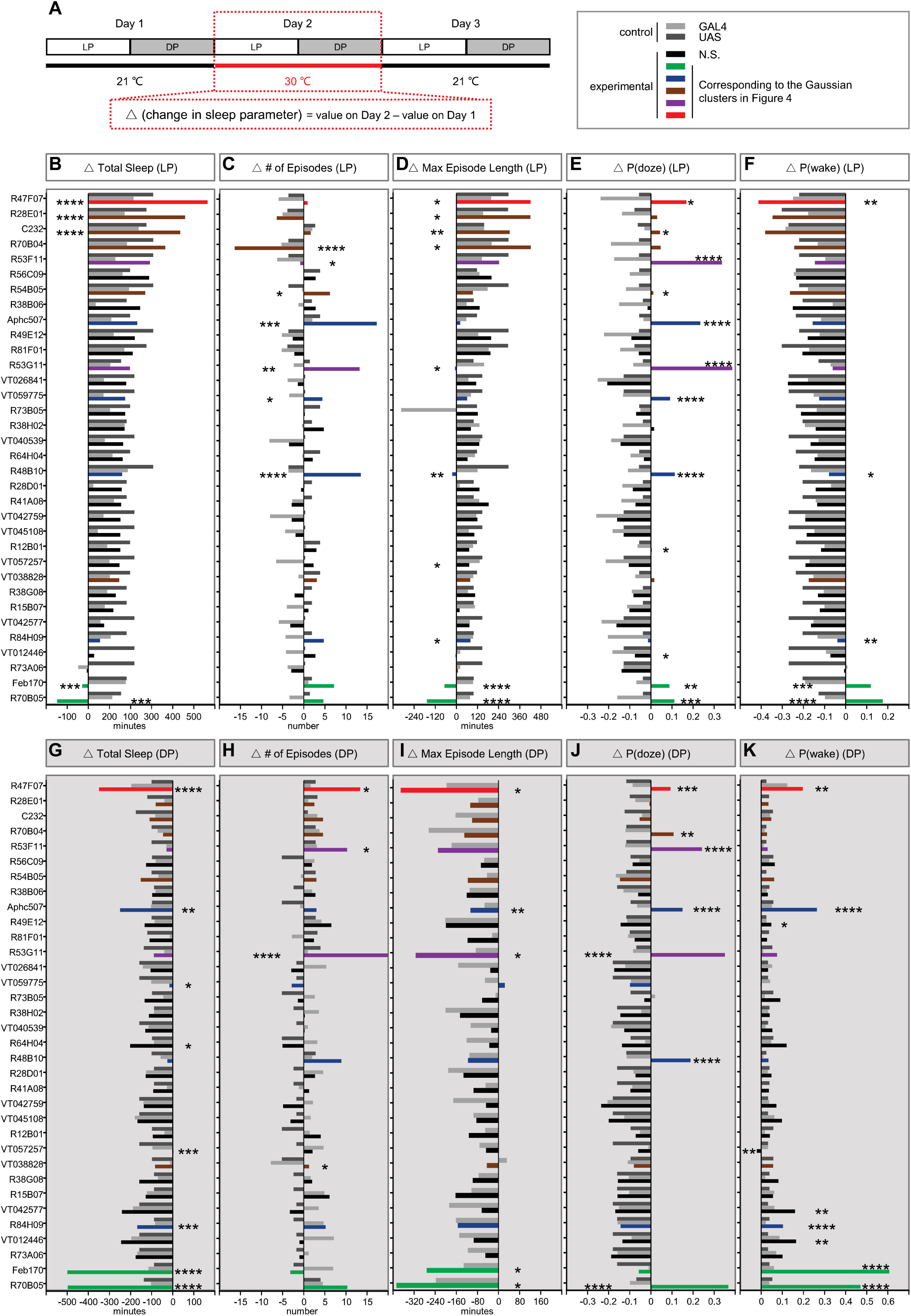
Sleep changes with activation of subtypes of ring neurons. (A) Design of the experiments and calculation of sleep parameters on the activation day (red dashed box). (B, G) Changes in sleep amount during day (LP) and night (DP). (C, H) Changes in number of sleep during daytime and night. (D, I) Changes in max sleep episodes during daytime and night. (E, J) Changes in P(doze) during daytime and night. (F, K) Changes in P(wake) during daytime and night. The colored and black bars represent the experimental groups. Color codes are consistent through all of the figures and are based on the daytime cluster analysis in Figure 4. The grey and dark grey bars indicate GAL4 control and UAS control, respectively. One-way ANOVA analysis and Dunn’s multiple comparisons test were used. Significance is marked by asterisks only when the experimental group is significantly different from both GAL4 and UAS controls, *p < 0.05; **p < 0.01; ***p < 0.001; ****p < 0.0001. Data are presented as mean ± SEM. LP, light period; DP, dark period.

**Figure 2.**
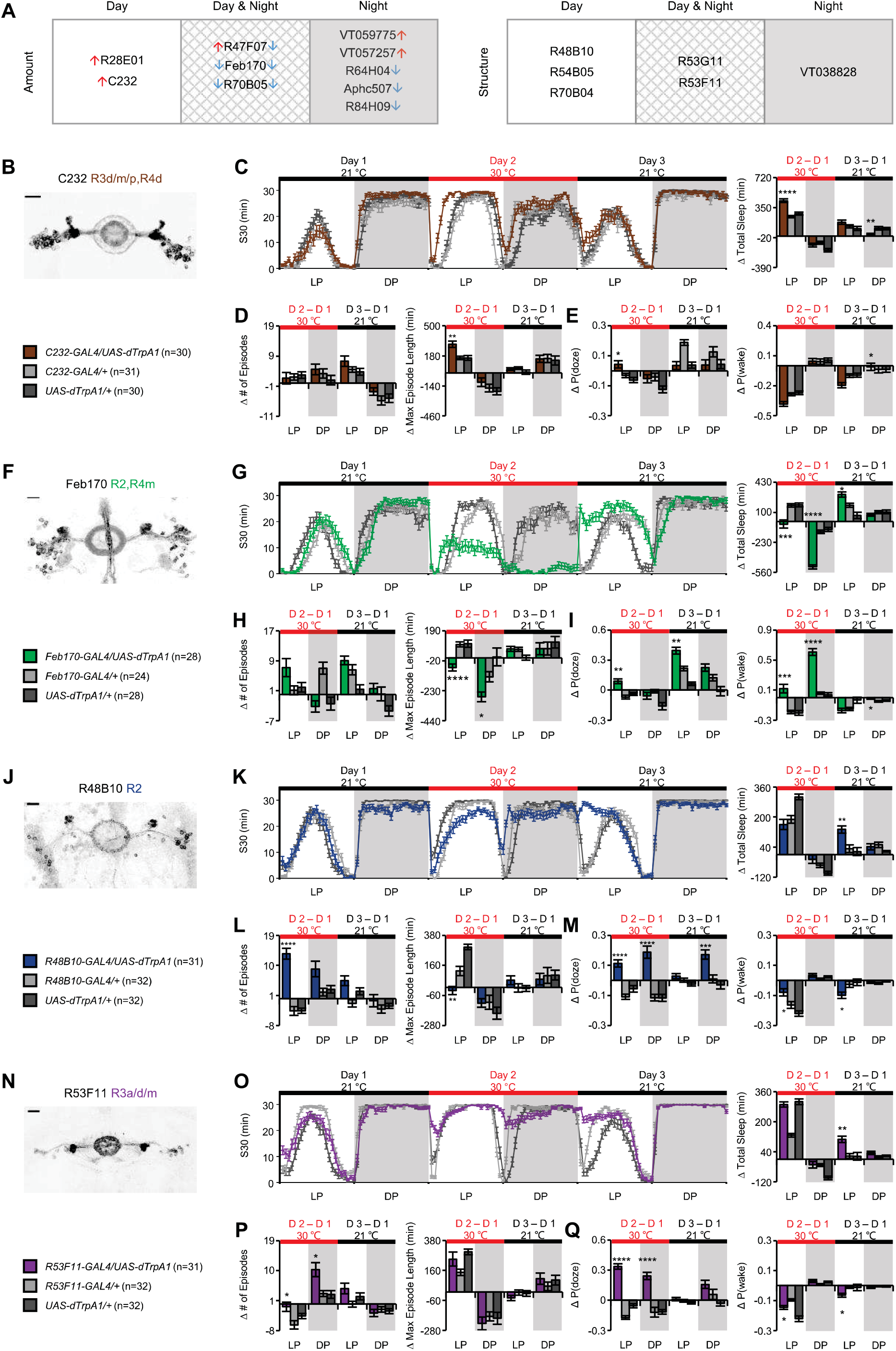
Complex effects on sleep homeostasis with thermoactivation of ring neurons. (A) Summary of the drivers exhibited significant changes in total amount of sleep and sleep structure, respectively. Arrows on the left and right represent changes during the day and night, respectively. Up arrows: increased total amount of sleep; down arrows: decreased total amount of sleep. Clusters represent the phenotypes observed day only, night only or both day and night. Expression patterns of c232-GAL4 (B), Feb170-GAL4 (F), R48B10-GAL4 (J) and R53F11-GAL4 (N). Sleep profiles with quantification of changes of sleep parameters of each driver: total sleep (C, G, K, O), the number of episodes and max episode length (D, H, L, P), and P(doze) and P(wake) (E, I, M, Q). Scale bar: 20 μm.

**Figure 3.**
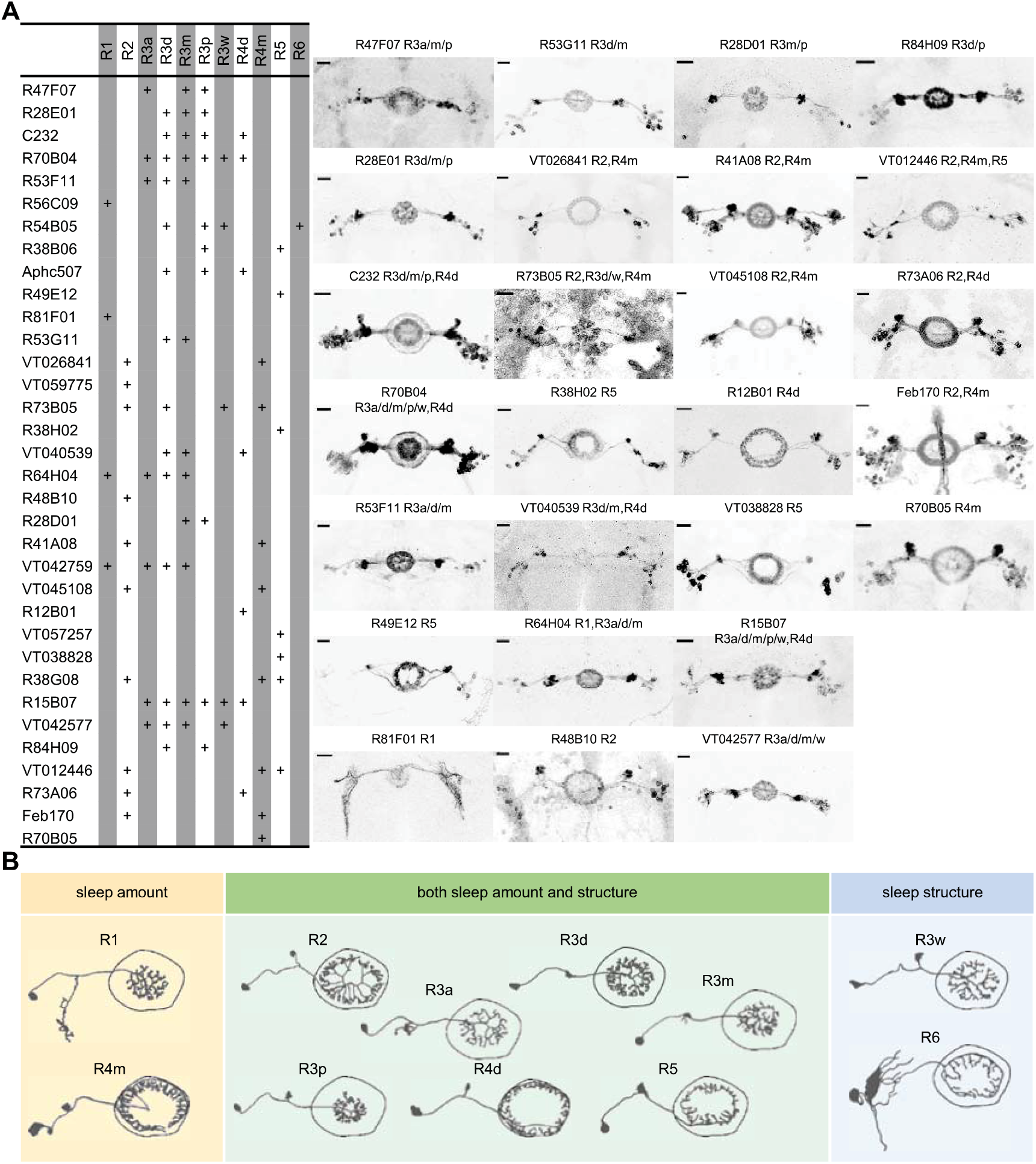
Expression patterns of EB drivers in the screen. Expression patterns of EB drivers in the screen. (A) Distinct subtypes of ring neurons were labeled by the 34 drivers. Expression patterns of all the publicly available drivers (26) are shown. (B) Single subtype of ring neurons that involved in the regulation of sleep amount (yellow), structure (blue), and both amount and structure (green).

**Figure 4.**
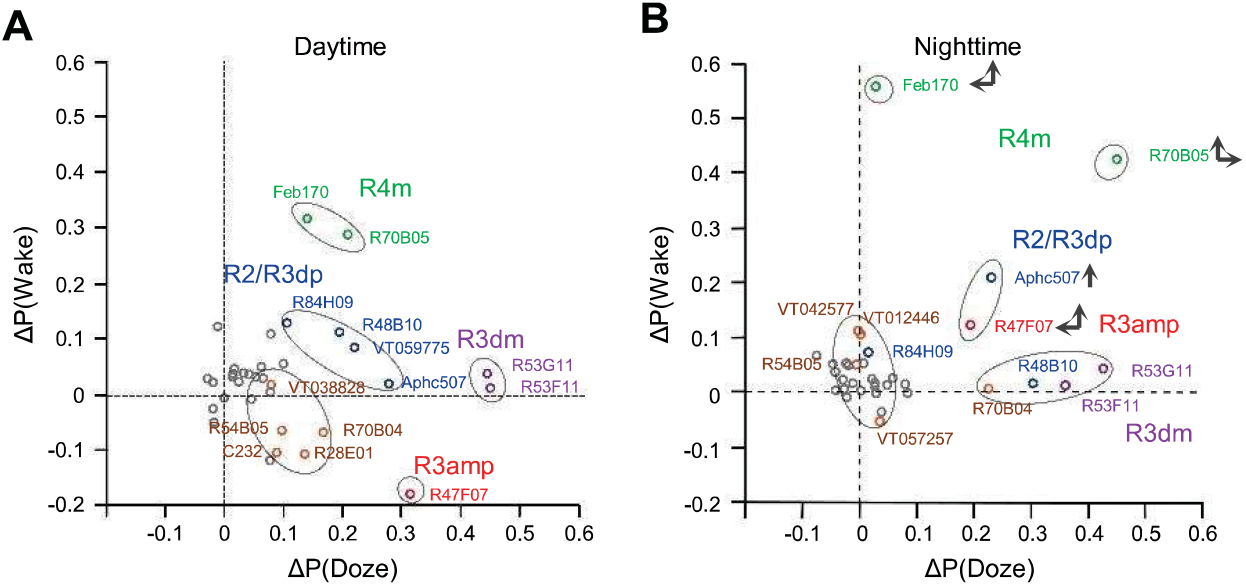
Association of changes in arousal and sleep drive with GAL4+ groups of ring neurons. Mixed Gaussian Model cluster analysis for drivers have similar patterns during the daytime (A) and at night (B). Grey dots: activation did not show significance in P(wake)/P(doze) analysis; green dots: increase in both P(wake) and P(doze); blue dots: mild increase in both P(wake) and P(doze); brown dots: weak increase in both P(wake) and P(doze); purple dots: increase only in P(doze); red dots: increase in P(doze) and decrease in P(wake). Vertical and horizontal arrows in nighttime panel represent shifts in location of P(doze) and P(wake) compared to the daytime.

**Figure 5.**
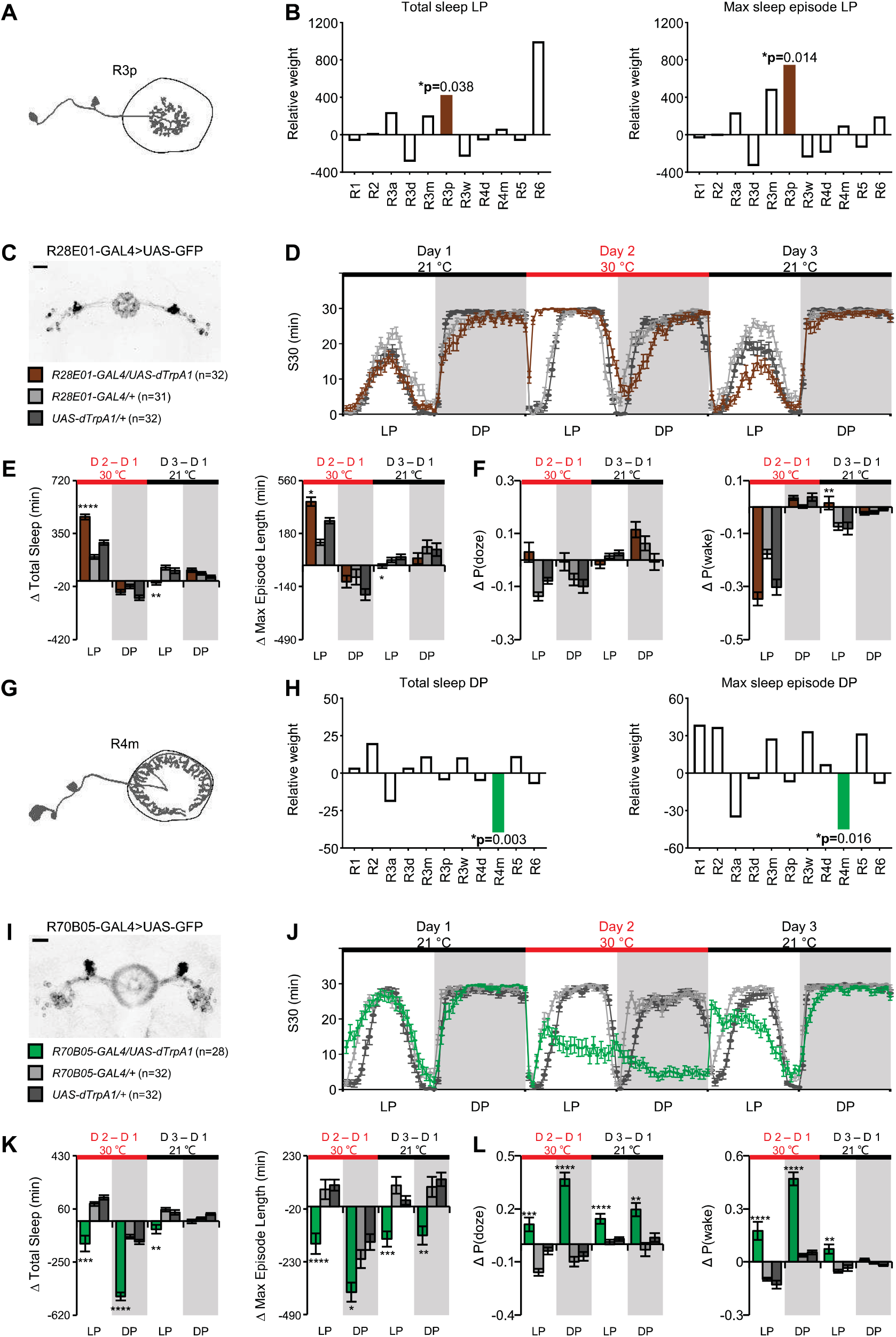
Two subtypes of ring neurons identified by general linear model that significantly contribute in the regulation of total sleep and episode length. (A, G) Schematic morphological pattern of a single R3p neuron and R4m neuron, respectively. (B, H) R3p neuron and R4m neuron are highly correlated to regulate daytime sleep and nighttime sleep, respectively. The weight of each subclasses was analyzed with a generalized linear model. (C, I) Expression pattern of R28E01-GAL4 and R70B05-GAL4 as representative for R3p and R4m. (D, J) Sleep profiles of total sleep before, during and after activation of R28E01-GAL4+ and R70B05-GAL4+ neurons with two controls. (E, K) Changes in total amount of sleep and max episodes on the activation day and the recovery day. R70B05-GAL4+ neurons not only significantly reduced nighttime sleep, also exhibited strong impact on reducing daytime sleep. (F) No detectable changes in P(doze) upon and after activation of R28E01-GAL4+ neurons. Weak elevation of P(wake) on the recovery day was found. (L) Strong increase in P(doze) were found when R70B05-GAL4+ neurons were activated, and this effect lasted with cessation of activation. Significant increase in P(wake) were also observed upon and after activation.

Activation of GAL4+ neurons produced many different patterns of change in the amount of sleep. During the daytime, a significant increase in total sleep was found when R47F07-GAL4+, R28E01-GAL4+, and C232-GAL4+ neurons were activated (Figure 1B). Since change in total sleep is often associated with change in sleep structure (Liu et al., 2019; Wiggin et al., 2020), we also evaluated the number of sleep episodes, episode length, and the behavioral transition probabilities, P(doze) and P(wake) (Wiggin et al., 2020) to further understand the changes in sleep drive and arousal threshold. The increased sleep observed in the above three drivers was accompanied by a significant increase in max episode length but no change in the number of episodes compared to their genetic controls (Figure 1C-D). These flies had increased P(doze) and decreased P(wake), suggesting that these neurons possibly contribute to increase sleep pressure and sleep depth (Figure 1E-F).

We also found cell groups which, when activated, induced a significant reduction in total sleep: Feb170-GAL4+ and R70B05-GAL4+ neurons (Figure 1B). Sleep reduction was associated with significant decreases in max episode length with no change in the number of episodes compared to their genetic controls (Figure 1C-D). The reduced sleep amount and episode length were possibly due to the increased P(doze) and P(wake) (Figure 1E-F), suggesting neurons labeled by these two drivers are involved in upregulation of sleep pressure and downregulation of sleep depth during the daytime.

Nighttime effects of thermogenetic neuron activation are more complex to interpret. Data have to be viewed in the context of the sleep-suppressing effects of elevated temperature on normal wild type animal sleep (Parisky et al., 2016; Jin et al., 2021). This temperature effect can be visualized in the continuous sleep plots for most of the GAL4 and UAS control lines in Figures 2 and 5 and Figures 2-1. VT059775-GAL4+ and VT057257-GAL4+ neuron activation led to almost no change of total sleep compared to their own baseline, but this reflects a significant difference from genetic controls, which respond to heat with at large reduction in sleep. These lines also had only small reductions in P(wake) compared to controls, implying that these neurons may be involved in sleep promotion by changing sleep depth (Figure 1G and 1K).

We also found a number of GAL4 drivers, including R47F07, Aphc507, R64H07, R84H07, Feb 170, and R70B05 which significantly reduced nighttime sleep amount compared to their controls, suggesting they contribute to promoting wakefulness (Figure 1G). These reductions in total sleep were accompanied by changes in sleep structure, featured as fragmentation where the number of episodes significantly increased and/or episode length reduced (Figure 1H-I). Many drivers exhibited increased P(doze) and P(wake) (Figure 1J-K), suggesting sleep pressure and sleep depth play important roles in nighttime sleep.

### Thermoactivation of ring neurons can change sleep structure independent of sleep amount

We also found cases where sleep structure was changed without alterations in total sleep, supporting the idea that structure can be regulated independently (Liu et al., 2019). Activation of neurons from several GAL4 drivers, including R70B04, R53F11, R54B05, R53G11, R48B10, and VT038828, resulted in significant change only in sleep structure. Except for R70B04, which induced consolidated daytime sleep with a decrease in the number of episodes and an increase in the episode length, all drivers mentioned above exhibited fragmented sleep either during the day or at night (Figure 1C-D and 1H-I). Fragmentation was accompanied by a robust increase in P(doze) for the majority drivers (Figure IE and J). P(doze) is believed to correlate with sleep pressure (Wiggin et al., 2020), suggesting the fragmentation reflects an increase in the probability of switching from wake to sleep i.e. high sleep drive.

The circadian period during which fragmentation occurred varied with GAL4 line. Daytime fragmentation was observed when R54B05-GAL4+ and R48B10-GAL4+ neurons were activated (Figure 1C), and nighttime fragmentation was seen when VT038828-GAL4+ neurons were activated (Figure 1H). Fragmentation of both day and night was found when R53G11-GAL4+ and R53F11-GAL4+ neurons were activated (Figure 1C-D and H-I).

The structural parameters that were altered were also variable. Three GAL4 drivers, R53F11, R54B05 and VT038828, only exhibited a significant increase in the number of episodes. R53G11 and R48B10 only showed reduced episode length. All of these changes contributed to increases in P(doze) with little or weak P(wake) effects, especially during the day (Figure 1E-F and J-K). Interestingly, R12B01-GAL4 did not exhibit detectable changes in the number of episodes or episode length, but had a significant increase in P(doze) compared to both controls (Figure 1E), suggesting a potential specific contribution of R28E01-GAL4+ neurons to control of sleep pressure. Taken together, changes in sleep structure are highly associated with P(doze), but when sleep structure changes are accompanied by changes in total sleep amount, P(wake) becomes an important component of the regulation.

### Thermoactivation of ring neurons has complex effects on sleep homeostasis

We summarized drivers with significant changes of total amount of sleep or sleep structure either during the day, at night or both (Figure 2A). We plotted sleep and changes in parameters over three days to provide a more nuanced picture of the lasting effects of activation of these neurons and present the lines ordered from largest to smallest rebound sleep on the recovery day (Figure 2B-Q, Figure 2-1). For some of the lines, the changes in total sleep appeared to activate homeostatic changes that were evident during the recovery day. Activation of C232-GAL4+ neurons, which increases sleep on the activation day, leads to a negative rebound (decrease in sleep) upon cessation of activation (Figure 2C). Activation of Feb170-GAL4+ neurons decreased sleep both in the day and night, and this was followed by a homeostatic rebound increase in sleep (Figure 2G). Activation of R48B10-GAL4+ or R53F11-GAL4+ neurons led to fragmentation during either the day or both in the day and night, and a robust homeostatic rebound increase occurred (Figures 2J-Q). Interestingly, some drivers exhibited decreased sleep without a rebound change in sleep afterwards, e.g. R64H04, R47F07 and R84H09 (Figure 2-1), suggesting that for these lines, sleep loss was either not able to be compensated for or was not “counted” by the homeostat. These may represent cell types that are not integrated into the homeostat (Seidner et al., 2015).

### Association of changes in arousal and sleep drive with GAL4+ groups of ring neurons

The majority of the GAL4 lines we screened contained more than one subtype of ring neuron (Figure 3A). To examine the linkage between ring neuron types and distinct aspects of sleep amount and/or sleep structure, we first separated drivers into two groups (Figure 2A): 1) those which exhibited changes in sleep amount; and 2) those which exhibited no change in sleep amount but had changes in sleep structure. Based on the time of day when the phenotype was observed (day only, night only or both day and night), we classed those drivers into three clusters. For lines that changed total sleep, we noted their effects in Figure 2A as increasing or decreasing. The second type of information we layered into the analysis was the identification of the subtypes of ring neurons in each line according to anatomical features and recent nomenclature (Omoto et al., 2018; Hulse et al., 2021) (Figure 3A). Based on this primary classification, many subtypes of ring neurons including R1, R2, R4m, R4d, R5 and many R3 subtypes (R3a, R3m, R3d, and R3p) may participate in the regulation of sleep amount (Figure 3B). Due to the multiplicity of ring neurons in these EB drivers, it was hard to *a priori* link a single subtype of EB neuron with a specific function in the regulation of sleep amount/structure. Thus, we employed statistical models to try to identify links between ring subtypes and phenotypes.

The first approach we used was aimed at determining the effects of the GAL4 lines (each of which has a different mixture of ring neuron subtypes) in regulating sleep. We used a mixed Gaussian model for changes in P(wake) or P(doze) on the activation day compared to the baseline day (Figure 4A-B). We chose to use these transition probabilities since they capture some of the more complex aspects of sleep: P(wake) correlates with arousal state/sleep depth, while P(doze) is a measure of sleep drive (Wiggin et al., 2020). A single value of ΔP(wake) and ΔP(doze) for each line was calculated by subtracting the average of the genetic controls for that driver (experimental ΔP – (UAS ΔP + GAL4 ΔP)/2)). These values were then plotted in ΔP(wake)-ΔP(doze) space and clustered with the model to find groups with similar effects on sleep depth and pressure. We identified 5 clusters of GAL4 lines for day and night, respectively (Figure 4A-B). These clusters define the color codes used in Figures 1, 2, 5 and Figure 2-1.

Using our anatomical analysis of these lines, we found that the lines within each cluster shared a common ring neuron subtype. During the daytime (Figure 4A), R4m (and perhaps R2 neurons) emerged as strong candidates for the regulation of sleep depth/arousal since they are present in lines that have high ΔP(wake) values. R3dm cells appeared to increase sleep drive; lines with these cells had large ΔP(doze). R2, R3d and R3p neurons were present in several clusters and did not appear to have unique functionality with regard to sleep depth and drive, but a role in facilitation of the effects of R2 and R3m neurons, or in more specialized functions in sleep structure, cannot be ruled out. We also observed that many drivers play different roles during the day and night (Figure 4B). For example, R70B05 exhibits relative strong P(wake) but weak P(doze) effect during the day, but at night increases its influence on P(doze); R47F07 has little effect on P(wake) in the day, but becomes much more wake-promoting at night.

### Association of specific ring neuron subtypes with changes in sleep parameters

Since the variable analyzed using Gaussian clustering was the GAL4 line, which is most often a collection of different ring neuron subtypes, the effects we saw could also be the result of particular combinations of subtypes rather than the result of one dominant subtype alone. To try to isolate effects specific to subtypes, and to look at more specific sleep parameters, we employed a second method to extract the contributions of each ring neuron subtype to functional outcomes. Using a general linear model (GLM) with ring neuron subtype as the variable allowed us to calculate the weights of the potential contribution of each subtype of ring neuron to all the sleep parameters for daytime and nighttime, respectively (Figure 5-1). R3p exhibited a significantly positive effect on daytime sleep amount which was associated with its positive weight in episode length (Figure 5A-B). As an example, activation of R28E01-GAL4+ neurons, which include the R3p subtype, elevated daytime sleep and max episode length (Figure 5C-E, Figure 5-1). But the R3p subtype had little effect on P(doze) or P(wake) (Figure 5F). We also found that R4m had a significantly inhibiting effect on total sleep at night and a negative effect on episode length (Figure 5G-H, Figure 5-1), consistent with the results of Gaussian clustering. R70B05-GAL4+ neurons include the R4m subtype, and activation of neurons labeled by this driver caused a dramatic reduction of sleep in both day and night which is likely due to the shortened episode length (Figure 5I-K). The effects of activation of these R70B05-GAL4+ neurons persisted into the recovery day, with flies exhibiting significantly elevated sleep pressure and lightened sleep depth (Figure 5L).

### Ring neuron synergy is important for sculpting sleep

Interestingly, there were effects uncovered in the GLM analysis that were not seen with GAL4 drivers that labelled only that specific subtype. R1 and R2 neurons exhibit a significantly negative weight in the number of episodes at night, suggesting that these neurons may contribute to consolidation of sleep structure (Figure 5-1). However, we failed to observe consolidation after activation of R1- or R2-specific GAL4 drivers; R56C09 and R81F01 had little significant effect on sleep structure (Figure 1), while activation of R48B10 produced a moderately strong increase in P(doze/wake) (Figure 2M). This suggests that the sleep consolidation effects of activating these neurons uncovered by the GLM requires co-activation of other subtypes.

Supporting the complexity of ring neuron subtype interactions, we observed that activation of the R47F07-GAL4 driver, which labels R3a, R3m and R3p ring neurons, induced increased daytime sleep but reduced nighttime sleep (Figure 4, Figure 2-1E-F). Increased daytime sleep was associated with an increase of episode length, explained by elevated sleep pressure and “deeper” sleep depth (Figure 4, Figure 2-1G-H). Opposite to the daytime change, reduced nighttime sleep was accompanied by fragmentation, resulting in increased sleep pressure and/or light sleep depth (Figure 4, Figure 2-1G-H). How these three subtypes of ring neurons coordinate to segregate, and effect a sign change on, day and night sleep still needs to be determined, but may provide insight into coordination of the EB circuit.

### Regulation of sleep fragmentation by a specific ring neuron subset

One of the interesting findings of this screen was that there appeared to be circuits which regulate sleep structure independent of sleep amount. These data were consistent with our previous studies which identified 5HT in EB as a modulator of sleep structure; activation of 5HT7-GAL4+ neurons fragmented sleep without changing the amount of sleep (Liu et al., 2019). 5HT7-GAL4+ neurons include R3d, R3p, and R4d subtypes (Hulse et al., 2021). To examine whether sleep structure regulation could be attributed to a specific subtype, we identified a driver R44D11-LexA that had an expression pattern similar to 5HT7-GAL4 (Figure 6A). LexA+ neurons overlapped nearly 79% with 5HT7-GAL4+ neurons (Figure 6C), but activation of R44D11-LexA+ neurons dis not induce sleep/structure changes upon activation (Figure 6-1). To test the hypothesis that sleep fragmentation might be induced by the non-overlapping population of 5HT7-GAL4+ neurons, we introduced LexAop-GAL80 to suppress the overlapping neurons between R44D11-LexA+ and 5HT7-GAL4+ neurons (Figure 6B). We found that activation of the non-overlapping 5HT7-GAL4+ neurons increased the number of episodes and reduced episode length (Figure 6D), suggesting that the non-overlapping neurons play a critical role in sleep fragmentation. Interestingly, the non-overlapping neurons morphologically are R3d subtypes (Figure 6B). This subtype of ring neuron was present in 4/6 of the lines we identified in this screen as affecting structure only (R70B04, R53F11, R54B05, R53G11) and there were also R3d neurons in some lines that fragmented sleep in addition to changing its amount (Aphc507, R84H09). The fact that not all lines which contain this ring neuron subtype fragment sleep may be due to interactions with other ring neuron types or heterogeneity within the R3d population.

**Figure 6.**
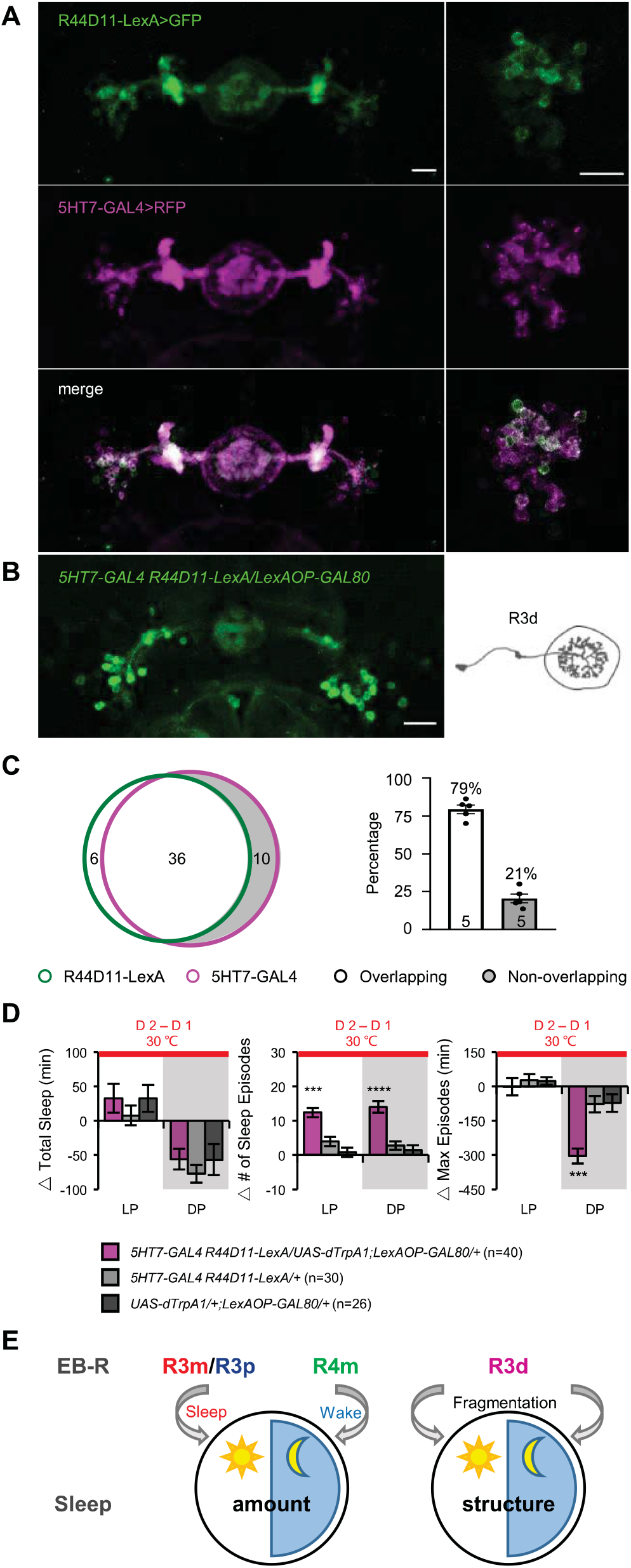
R3d neurons contribute to sleep fragmentation. (A) Expression pattern of R44D11-LexA+ neurons and 5HT7-GAL4+ neurons labeled by GFP and REP, respectively. 79% of the R44D11-LexA+ neurons (green) overlap with 5HT7-GAL4+ neurons (magenta). (B) R3d populations are labeled by suppressing the overlapped neurons of R44D11-LexA+ and 5HT7-GAL4+ by using LexAOP-GAL80. Bar: 20 μm. (C) Venn diagram shows the overlapping and non-overlapping cells between 5HT7-GAL4+ and R44D1 l-LexA+. Bar graph represents the quantified ratio of non-overlapping neurons. (D) Activation the non-overlapping R3d neurons fragments sleep without significant effect on total amount during LP and DP. (E) Schematic of sleep/structure regulation by multiple subtypes of ring neurons.

## DISCUSSION

Sleep is crucial for survival and overall health across animal kingdoms. Fly sleep exhibits the majority of the highly conserved features of vertebrate sleep, and the tractability of *Drosophila* as an experimental model has produced a growing number of studies which contribute to our knowledge of sleep mechanisms and circuits. Besides the importance in learning and memory of the mushroom body (MB), multiple subtypes of intrinsic MB Kenyon cells (KCs) have been identified for the control of sleep (Joiner et al., 2006; Sitaraman et al., 2015; Artiushin and Sehgal, 2017; Bringmann, 2018). For example, α’β’ and γm KCs contribute to wake promotion, and γd KCs contribute to sleep promotion (Sitaraman et al., 2015). A pair of GABAergic and serotonergic dorsal paired medial (DPM) neurons, which are MB extrinsic projecting neurons and play a role in memory consolidation (Keene et al., 2004; Keene et al., 2006; Zhang et al., 2013), were shown to be involved in sleep promotion (Haynes et al., 2015). Dopaminergic PPL1 and PPM3 neurons that project to different layers of fan-shaped body (FB) have been shown their specific roles in promoting wake, via suppression of the FB which is thought as a sleep induction center (Liu et al., 2012; Ueno et al., 2012; Pimentel et al., 2016).

Many of these brain structures have been implicated in multiple behaviors. Like the MB and FB mentioned above, the EB has been shown to integrate sensory inputs to formulate locomotor output commands, but understanding of its role in sleep is still limited. In the present study, we identified subtypes of ring neurons that regulate sleep/structure by: 1) screening a small collection of EB drivers using thermogenetic activation; and 2) specifying the roles of several single subtypes in different sleep components employing two models and intersection strategies. We found that R3m/R3p neurons contribute to daytime sleep, R4m neurons to wakefulness, and R3d neurons fragment sleep structure (Figure 6E).

The role of these neurons in sleep may be intimately involved with their other functions. Previous studies found that R2, R3, R4d and R4m subtypes appear to be tuned to visual stimuli (Shiozaki and Kazama, 2017; Fisher et al., 2019; Kim et al., 2019; Hardcastle et al., 2021). This sensory input may be an important cue to change sleep/wake status, and is likely influenced by the circadian system. Previous study showed that the R5 subtype is linked to the control of sleep homeostasis and stabilization of sleep structure (Liu et al., 2016; Liu et al., 2019), and our analysis support previous findings. A recent study released on bioRxiv identified two subtypes, sleep promoting R3m neurons and wake promoting R3d neurons (Aleman et al., 2021). Consistently, we also observed that R3m contributes both sleep amount and sleep structure. 5HT7-GAL4+ neurons play an important role in sleep maintenance, when they are activated, sleep became fragmented (Liu et al., 2019). According to a recent anatomical analysis (Hulse et al., 2021), 5HT7-GAL4+ neurons include R3d, R3p and R4d subtypes, and we narrowed the fragmentation effect down to a specific subtype, R3d in the present study. However, more efforts are still needed to understand how a certain subtype of ring neuron responds to sensory inputs and how neuronal activity patterns form in the network. Future work examining the neural activity of each subtype of ring neurons that control distinct sleep components and the interaction with other behaviors may reveal fundamental information about the rules of the coding and integration of the brain.

## Acknowledgements

This work is supported by National Natural Science Foundation of China (32071009 to CL), Guangdong Basic and Applied Basic Research Foundation (2020A1515011055 to CL), CAS Key Laboratory of Brain Connectome and Manipulation (2019DP173024), National Institutes of Health grant R01MH67284 to LCG.

---

**Figure 2-1.**
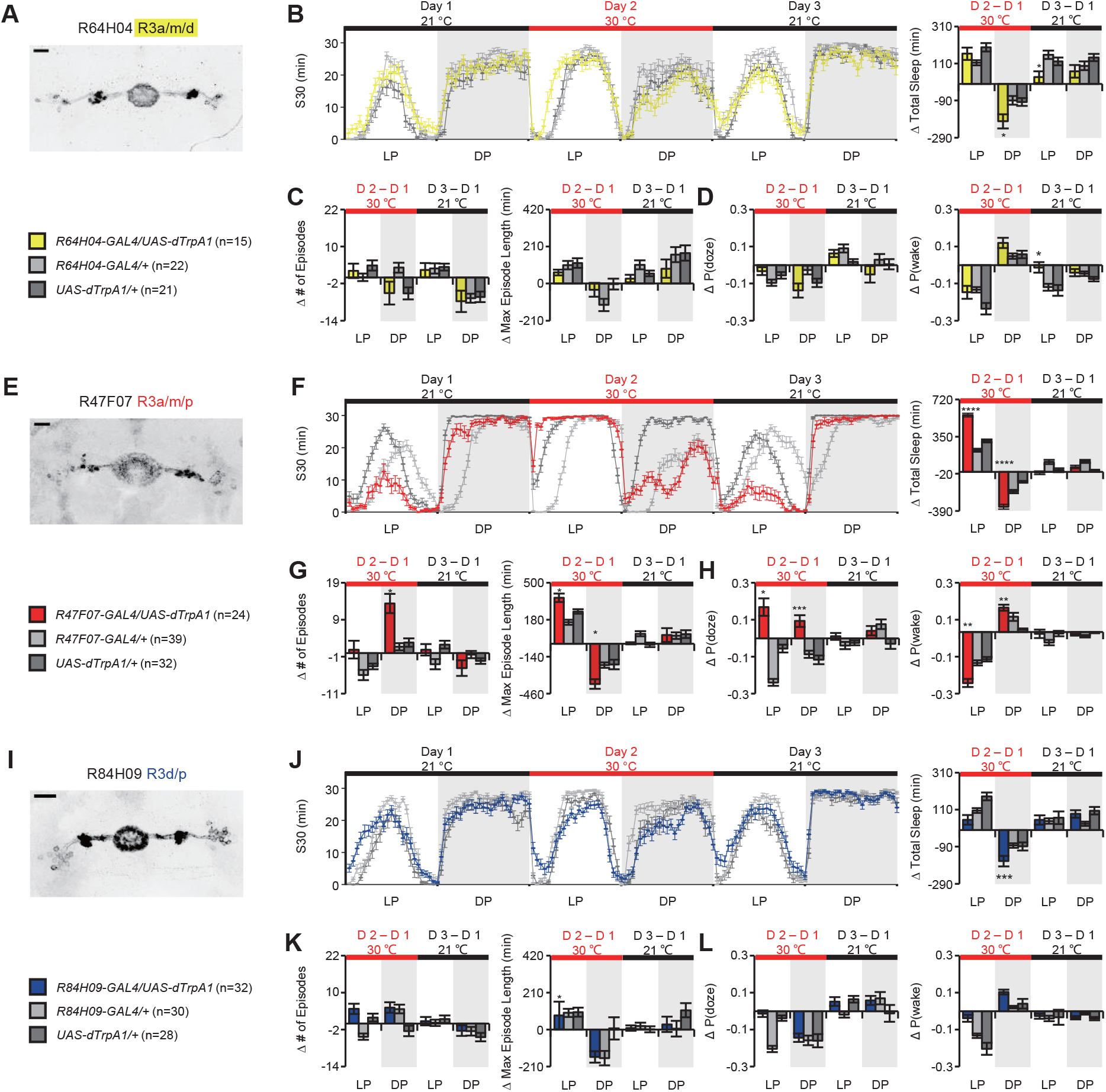
(A-D) Decreased nighttime sleep often fails to induce rebound sleep upon cessation of thermoactivation of ring neurons. (A) Expression pattern of R64HO4-GAL4 which labels R3a, R3m and R3d neurons. (B) Sleep profile and quantification of total sleep before, during and after activation. Activation of R64H04-GAL4+ neurons reduced total sleep at night, and a persisting reduced sleep upon cessation of activation. (C) No significant change was observed in the number of episodes and max episode length. (D) No change of P(doze) was observed, and significantly higher change in P(wake) than controls was found upon cessation of activation. **(E-L) Drivers involved in the regulation of sleep amount and/or structure do not exhibit homeostatic rebound upon cessation of thermoactivation**. Expression pattern, sleep profile, quantification of sleep amount, sleep structure, and sleep drive/arousal threshold of each driver were presented. (E-H) R47F07-GAL4. (I-L) R84H09-GAL4.

**Figure 2-2.**
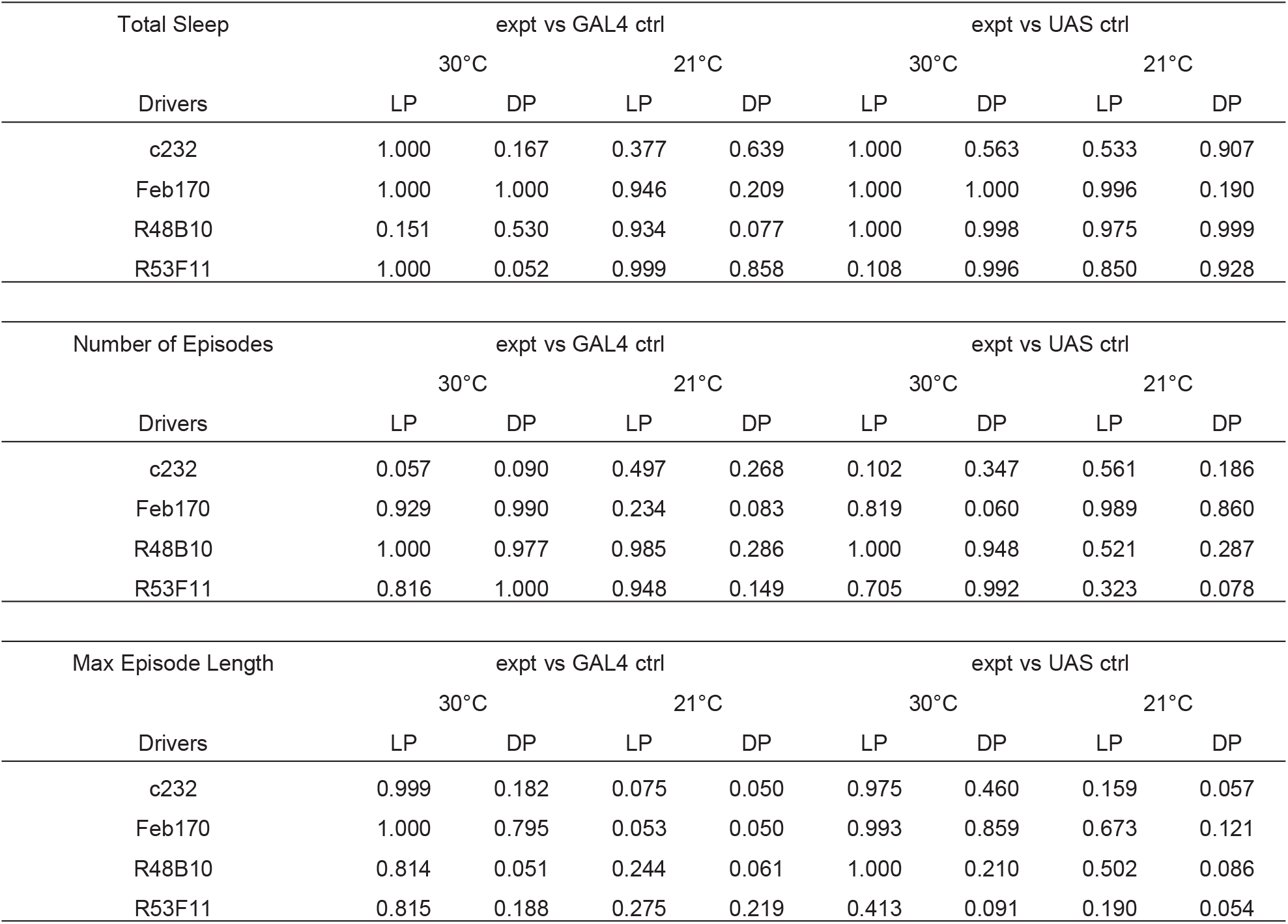
Power analysis for the sample size of drivers employed in Figure 2.

**Figure 5-1.**
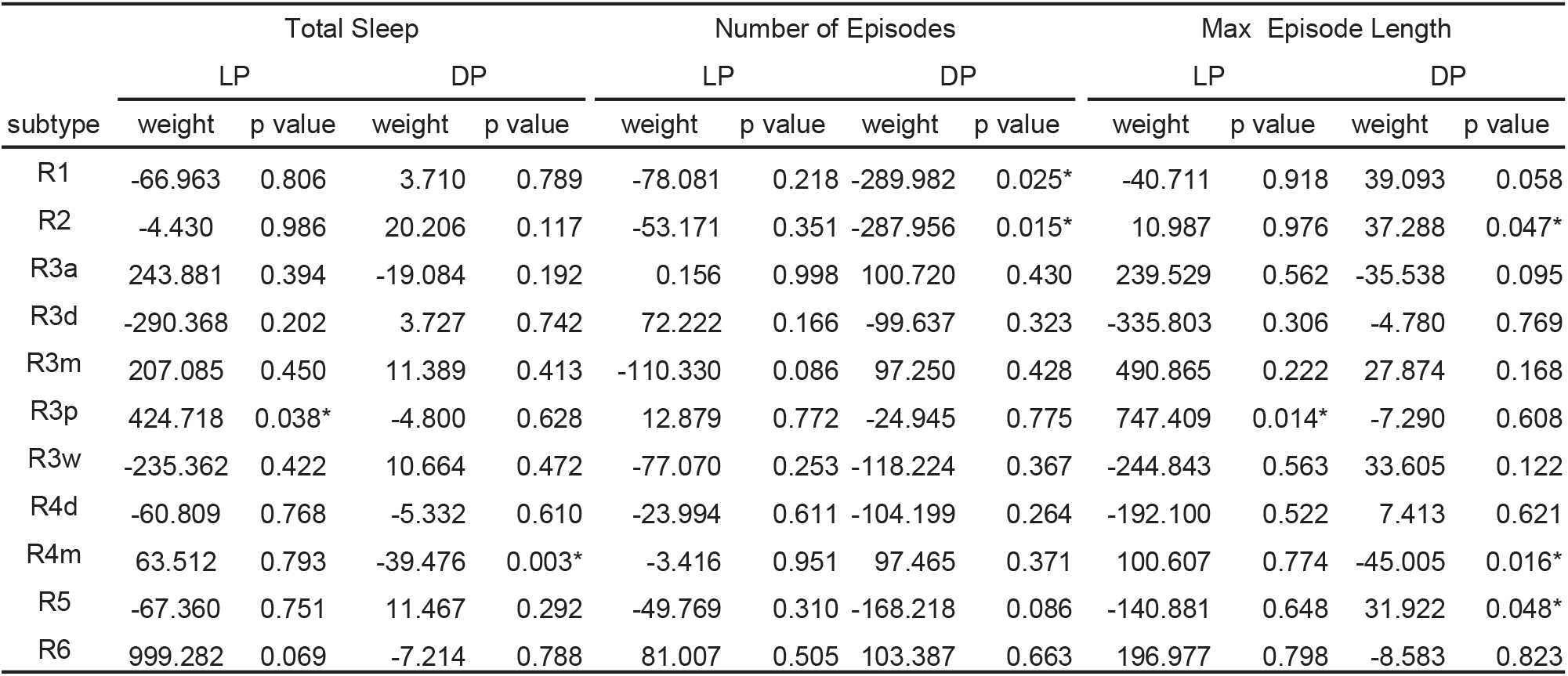
Statistical table of the weight of the effect of subclasses of ring neurons on the sleep using a generalized linear model. * p<0.05.

**Figure 5-2.**
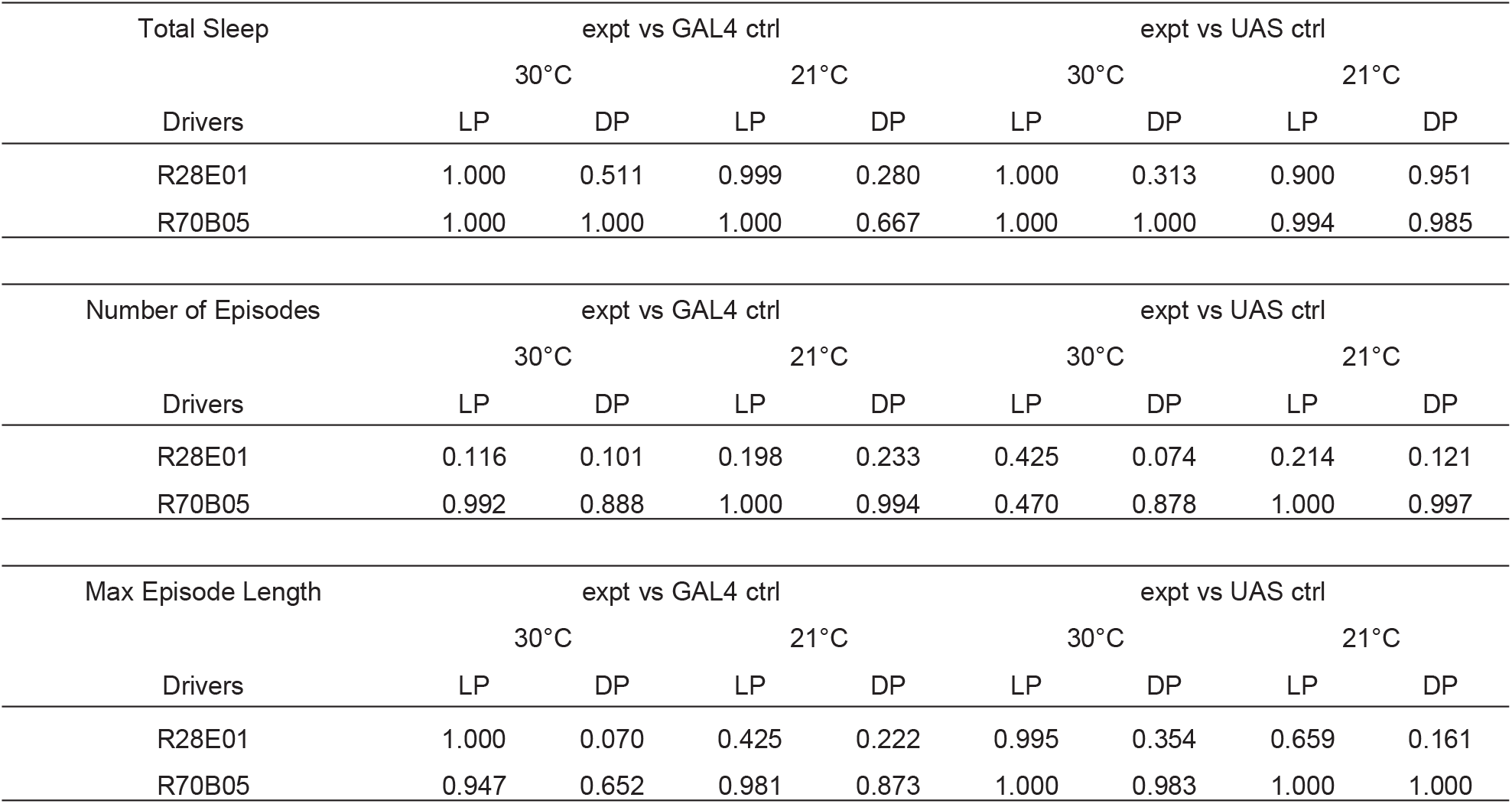
Power analysis for the sample size of drivers employed in Figure 5.

**Figure 6-1.**
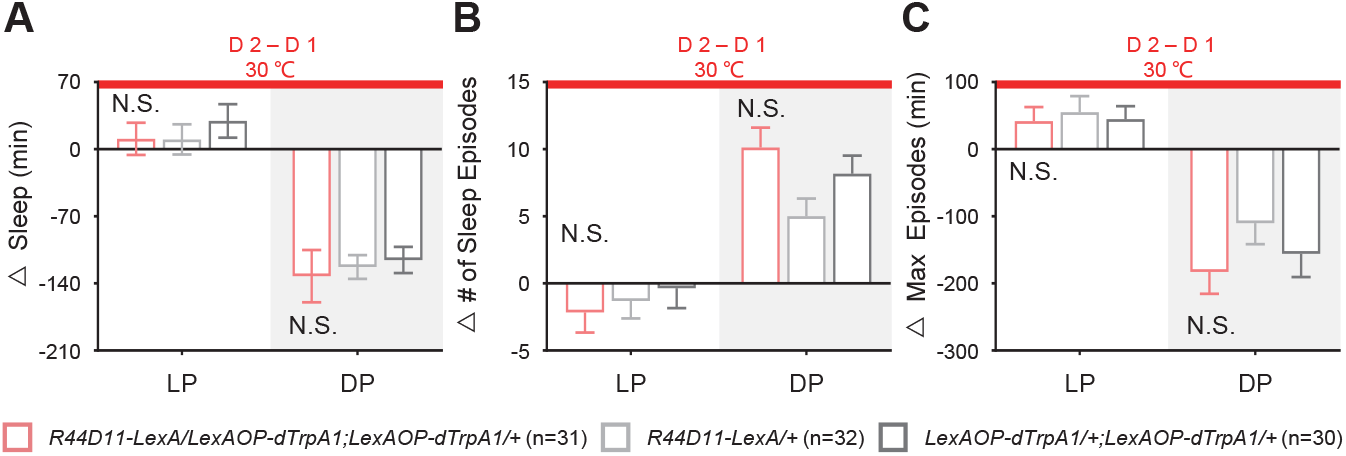
Activation of R44D11-LexA+ neurons do not change sleep or sleep structure. Upon the activation, no significant change was detected compared to both genetic controls in total sleep (A), the number of episodes (B), and max episode length (C) during both day and night.

